# The crystal structure of Nictaba reveals its carbohydrate-binding properties and a new lectin dimerization mode

**DOI:** 10.1101/2024.05.06.592670

**Authors:** Yehudi Bloch, Vinicius J.S. Osterne, Savvas N. Savvides, Els J.M. Van Damme

## Abstract

Nictaba is a (GlcNAc)n-binding, stress-inducible lectin from Nicotiana tabacum that serves as a representative for the family of Nictaba-related lectins, a group of proteins that play pivotal roles in plant defense mechanisms and stress response pathways. Despite extensive research into the biological activities and physiological role(s) of the lectin, the three-dimensional structure of Nictaba remained largely unknown. Here, we report crystal structures for Nictaba in the apo form and bound to chitotriose. The structures reveal a jelly-roll fold for the Nictaba protomer similar to the cucumber Cus17 lectin, but show an unprecedented dimerization mode among all structurally characterized lectins to date. The chitotriose binding mode, similar to Cus17, centers around the central GlcNAc residue, providing insights into the determinants of specificity of Nictaba towards carbohydrate structures. By integrating these structural insights with inputs from glycan arrays, molecular docking, and molecular dynamics simulations, we propose that Nictaba employs a single carbohydrate-recognition domain within each of the two subunits in the dimer to display pronounced specificity towards GlcNAc-containing carbohydrates. Furthermore, we identified amino acid residues involved in the extended binding site capable of accommodating structurally diverse high-mannose and complex N-glycans. Glycan array and in silico analyses revealed interactions centered around the conserved Man3GlcNAc2 core, explaining the broad recognition of N-glycan structures. Collectively, the structural and biochemical insights presented here fill a hitherto substantial void into the atlas of lectin structure-function relationships and pave the way for future developments in plant stress biology and lectin-based applications.

## 1. INTRODUCTION

Evidence accumulating over the last decades clearly showed that protein-carbohydrate interactions are key in a multitude of physiological processes across all kingdoms of life. With more and more protein-carbohydrate interactions being studied their importance and involvement in biochemical processes and signaling cascades has come to the surface, and the ‘glycocode’ has been defined. The sugar code or glycocode refers to the information present in the sugar structures on glycoconjugates (Wisnovsky & Bertozzi, 2022). For instance, glycans are known to have important roles in immune responses, developmental processes, pathogen recognition, stress responses, among others (Lin et al., 2020; Strasser, 2022; Varki, 2017) and therefore glycan signatures are considered as a checkpoint for biological processes (Seeberger et al., 2024; Zhou et al., 2018).

Although glycans are involved in a wide range of biological processes ranging from embryogenesis to protein trafficking and pathogen infection, our understanding of the glycocode remains incomplete. In this regard, structural and molecular insights into the repertoire of protein-carbohydrate structure-function landscapes are critical for the development of molecular tools towards deciphering the glycocode (Ambrosi et al., 2005; Wisnovsky & Bertozzi, 2022). Among the carbohydrate-binding proteins, lectins have received special interest since lectin domains can specifically recognize and bind to sugar moieties in a reversible way (Peumans & Van Damme, 1995), and, for this reason, these proteins are recognized as the main glycocode-decoders. Thus, a more complete understanding of lectin-carbohydrate interactions is desirable.

Plant lectins have been the subject of detailed studies for more than 130 years. In the early days of lectinology, their study focused mainly on the characterization of new lectins from different plant sources, but nowadays the attention has shifted to more functional analyses aiming to understand the role of lectins in protein-carbohydrate interactions important for plant growth and development (Tsaneva & Van Damme, 2020). At present the large and heterogeneous group of plant lectins can be classified in 12 families of lectins based on the sequence of the lectin domain, resulting in a classification of lectins based on structural characteristics rather than carbohydrate-binding specificities (Van Damme et al., 2008). Indeed, lectin domains from different families can recognize and bind similar carbohydrate structures. Though original studies focused on the interaction of lectins with monosaccharides, functional studies highlighted that the natural ligands for lectins are more likely complex glycan structures (Mattox & Bailey-Kellogg, 2021).

The *Nictotiana tabacum* agglutinin, abbreviated as Nictaba was first reported in 2002 as a dimeric 38 kDa protein that was expressed in tobacco leaves after treatment with the methyl ester of the plant hormone jasmonate, and was one of the first so-called stress-inducible plant lectins to be discovered (Chen et al., 2002). Further research revealed that lectin expression is also regulated by cold and insect herbivory (Delporte et al., 2013; Vandenborre et al., 2009). Based on hapten inhibition assays Nictaba was originally described as a (GlcNAc)_n_ binding lectin. The interaction with GlcNAc oligomers was confirmed by surface plasmon resonance experiments and showed best interaction with chitotriose (Chen et al., 2002). Detailed binding studies using glycan arrays also revealed Nictaba interaction with multiple high-mannose as well as complex *N*-glycans (Lannoo et al., 2006).

Nictaba resides in the cytoplasm but is also found within the plant cell nucleus, where it interacts with histone proteins modified by O-GlcNAc. As a result, the lectin is thought to function as a regulator of gene transcription (Schouppe et al., 2011). Overexpression of Nictaba and certain Nictaba homologues from *Arabidopsis* or soybean in transgenic plants revealed the importance of these lectins in the stress responses of plants to bacterial infection, nematodes and insect attack (Eggermont et al., 2017; Van Holle et al., 2016; Wojszko et al., 2023).

Since Nictaba-related sequences have been retrieved from genomes of many flowering plants Nictaba is considered the prototype of a widespread family of lectins. Screening of genome and transcriptome databases further revealed that the Nictaba domain can occur both as a standalone domain but also often as part of a chimeric protein in which the Nictaba domain is fused to other domains, such as the F-box or TIR domain (Delporte et al., 2015; Van Damme et al., 2004), suggesting that throughout evolution the carbohydrate-binding Nictaba domain has been used as a building block to create multidomain proteins.

BLAST sequence alignments highlighted that the Nictaba sequence is evolutionarily related to the so-called phloem protein 2 (PP2) lectins. In 1978, Sabnis and Hart first reported that one of the most abundant proteins in the phloem of *Cucurbita maxima* was a lectin (Sabnis & Hart, 1979). The lectin from *Cucurbita pepo* fruit was purified by affinity chromatography using chitin oligosaccharides (Allen, 1979). Since then, numerous lectins termed PP2 lectins, have been characterized mainly from Cucurbitaceae species (Dinant et al., 2003; Swamy et al., 2022). Sequence alignments revealed that the Nictaba sequence is missing the N-terminal sequence (50-80 amino acid residues) and the cysteine-rich C-terminal sequence (5 amino acids) that are present in 24-26 kDa PP2 lectins (Lannoo & Van Damme, 2010; Van Damme et al., 2004). Some Cucurbitaceae species contain a second PP2-type lectin domain of ∼17 kDa in which these additional domains are lacking (Swamy et al., 2022; Swamy & Mondal, 2023). Similar to Nictaba, all PP2 lectins share a carbohydrate-recognition domain that preferentially interacts with chitin, chito-oligosaccharides and *N*-glycans. However, unlike PP2 lectins that are exclusively and continuously expressed in the companion cells of the phloem and then translocated into the phloem sap, Nictaba is a stress-inducible protein residing in the nucleus and the cytoplasmic compartment, it is not present in the phloem (Balachandran et al., 1997; Lannoo & Van Damme, 2010). Although Nictaba and PP2 lectins share a similar carbohydrate-recognition domain they probably fulfill different biological roles in the plant.

Because of their carbohydrate binding activity plant lectins have evolved as tools for glycobiology research. For instance, several lectins have been used to understand the interaction between viruses and their target cells, since the infection process mainly relies on protein-carbohydrate interactions from target cells with glycan structures on the surface of enveloped viruses (Nabi-Afjadi et al., 2022; Santisteban Celis & Matoba, 2024). Studies aiming at the screening of a panel of lectins for antiviral activity revealed that Nictaba showed antiviral activity against human immunodeficiency virus I as well as the coronavirus SARS-CoV (Gordts et al., 2015; Keyaerts et al., 2007).

Though the three-dimensional structure of the lectin domain for 10 out of 12 lectin families has been resolved, structural data on Nictaba-related lectins are lacking, with the only experimental structure for the phloem lectin from *Cucumis sativus* (Cus17) (Bobbili et al., 2023). To address this gap, this study aimed to elucidate the structure of Nictaba. Recombinant Nictaba was purified from *Escherichia coli* and structurally characterized by X-ray crystallography. Additionally, the carbohydrate-binding properties of the lectin were unveiled through a combination of structural insights derived from a complex of Nictaba with chitotriose, molecular docking, and molecular dynamics. The obtained data represent a leap forward in the understanding of stress-inducible lectins, the specific function of Nictaba and Nictaba-related lectins, and possible new biological applications for this lectin.

## 2. RESULTS

### 2.1 Nictaba folds as a jelly roll and adopts a new lectin dimerization mode

To enable structural studies of Nictaba by X-ray crystallography, we expressed Nictaba in *E. coli* and purified monodisperse preparations of recombinant Nictaba with and without a (His)_6_-tag (**Supplementary Figure 1**). Intriguingly and in contrast to previous reports, recombinant Nictaba was found to be a tetrameric protein in solution (75.6 kDa ± 0.5 kDa compared to the expected MW = 4 x 19.1 kDa = 76.4 kDa for a tetramer) by Size-exclusion Chromatography in line with Multi-angle Laser Light Scattering (SEC-MALLS). Recombinant untagged Nictaba was propagated to crystallization trials and led to diffraction quality crystals that yielded X-ray diffraction data to 2.15 Å resolution using synchrotron X-rays (**Table 1**). The crystal structure of Nictaba was determined by Single-wavelength Anomalous Diffraction (SAD) (**Supplementary Figure 2**) and revealed 8 copies of Nictaba subunits in the asymmetric unit of the crystal organized as 4 sets of dimers each obeying a C2 symmetry. Tetrameric assemblies that would be consistent with our SEC-MALLS data of purified Nictaba were not unambiguously observed. The Nictaba subunit adopts a jelly-roll with an ancillary beta hairpin (annotated as βB’ and βB’’) originating from the B β-strand at the edge of the domain (**Figure 1A,B**). The tetralysine motif Lys102-105, which probably acts as a nuclear localization signal, is part of a structured loop which is solvent accessible. Despite the similarity between the folds of Nictaba and legume lectins, Nictaba is not a metalloprotein and does not contain any non-proline *cis*-peptides in its structure.

**Table 1.** X-ray diffraction data statistics and model refinement statistics * Friedel pairs assumed not equivalent.

|  | Nictaba Apo | I soak* | Eu soak* | Nictaba NAG3 |
| --- | --- | --- | --- | --- |
| **PDB code** | 8qmg | 8qmg (data only) | 8qmg (data only) | 8ad2 |
| **Dataset statistics** |  |  |  |  |
| Condition | 91 mM ZnAc, 91 mM ZnCl<sub>2</sub>, 91 mM Bis-Tris pH 7.5, and 18.2% Medium PEG smear | BCS E3 Opt B12 2P+1ML |  | 91 mM ZnAc, 91 mM ZnCl<sub>2</sub>, 91 mM Bis-Tris pH 7.5, and 18.2% Medium PEG smear |
| Cryo | 17% ZW221 | 17% ZW221 + 400 mM NaI | 17% ZW221 + 100 mM Eu DO3A | 5% DMSO, 5% EtGlyc, 5% Glycerol+1mM NAG3 |
| Unit Cell Parameters (Å) | 66.51 145.97 82.05 | 69.47 145.71 82.08 | 70.16 151.43 82.52 | 66.66 146.16 81.91 |
| (°) | 90.00 95.95 90.00 | 90.00 96.74 90.00 | 90.000 97.632 90.000 | 90.00 95.87 90.00 |
| Space group | P 2<sub>1</sub> (n°4) | P 2<sub>1</sub> (n°4) | P 2<sub>1</sub> (n°4) | P 2<sub>1</sub> (n°4) |
| Wavelength (Å) | 1.0332 | 2.0664 | 1.5326 | 0.9763 |
| Resolution (Å) | 81.61-2.15 (2.28-2.15) | 81.24-2.50 (2.57-2.50) | 69.54 – 2.60 (2.67-2.60) | 60.39-2.00 (2.12-2.00) |
| Reflection observed | 509983 (42289) | 1829977 (102282) | 2735371 (144006) | 397928 (42319) |
| unique | 72679 (6603) | 108471 (7800) | 101535 (6281) | 97998 (11697) |
| multiplicity | 7.02 (6.40) | 16.87 (13.11) | 26.94 (22.93) | 4.06 (3.62) |
| Completeness (%) | 86.1 (48.5) | 98.2 (94.7) | 98.1 (82.9) | 93.4 (69.2) |
| I/σ(I) | 7.93 (1.38) | 8.96 (0.60) | 11.03 (1.11) | 8.01 (1.23) |
| R-meas (%) | 21.9 (114.3) | 26.8 (348.0) | 28.2 (213.6) | 13.2 (110.9) |
| CC(1/2) | 99.2 (54.4) | 99.6 (17.3) | 99.7 (48.1) | 99.6 (51.0) |
| SigAno |  | 1.34 (0.59) | 1.268 (0.612) |  |
| Wilson B (Å<sup>2</sup>) | 35.12 | 59.79 | 55.61 | 38.34 |
| **Refinement Statistics** |  |  |  |  |
| Reflections used in refinement | 65010 (4657) |  |  | 97984 (6718) |

| Reflections used<br>for R-free | 2274 (163) |  |  | 2000 (138) |
| --- | --- | --- | --- | --- |
| R-work | 0.2016 (0.2743) |  |  | 0.1983 (0.3174) |
| R-free | 0.2387 (0.3290) |  |  | 0.2284 (0.3516) |
| Number of non-<br>hydrogen atoms | 11704 |  |  | 11698 |
| macromolecules | 10788 |  |  | 10792 |
| ligands | 24 |  |  | 196 |
| solvent | 892 |  |  | 710 |
| Protein residues | 1319 |  |  | 1319 |
| RMS(bonds) (Å) | 0.005 |  |  | 0.013 |
| RMS(angles) (°) | 0.54 |  |  | 1.24 |
| Ramachandran<br>favored (%) | 96.47 |  |  | 96.85 |
| allowed (%) | 3.07 |  |  | 2.76 |
| outliers (%) | 0.46 |  |  | 0.38 |
| Rotamer outliers<br>(%) | 1.14 |  |  | 1.15 |
| Clashscore | 3.42 |  |  | 4.34 |
| Average B-factor<br>(Å²) | 38.85 |  |  | 38.86 |
| macromolecules | 38.79 |  |  | 38.39 |
| ligands | 62.91 |  |  | 60.17 |
| solvent | 38.89 |  |  | 40.07 |

**Figure 1.**
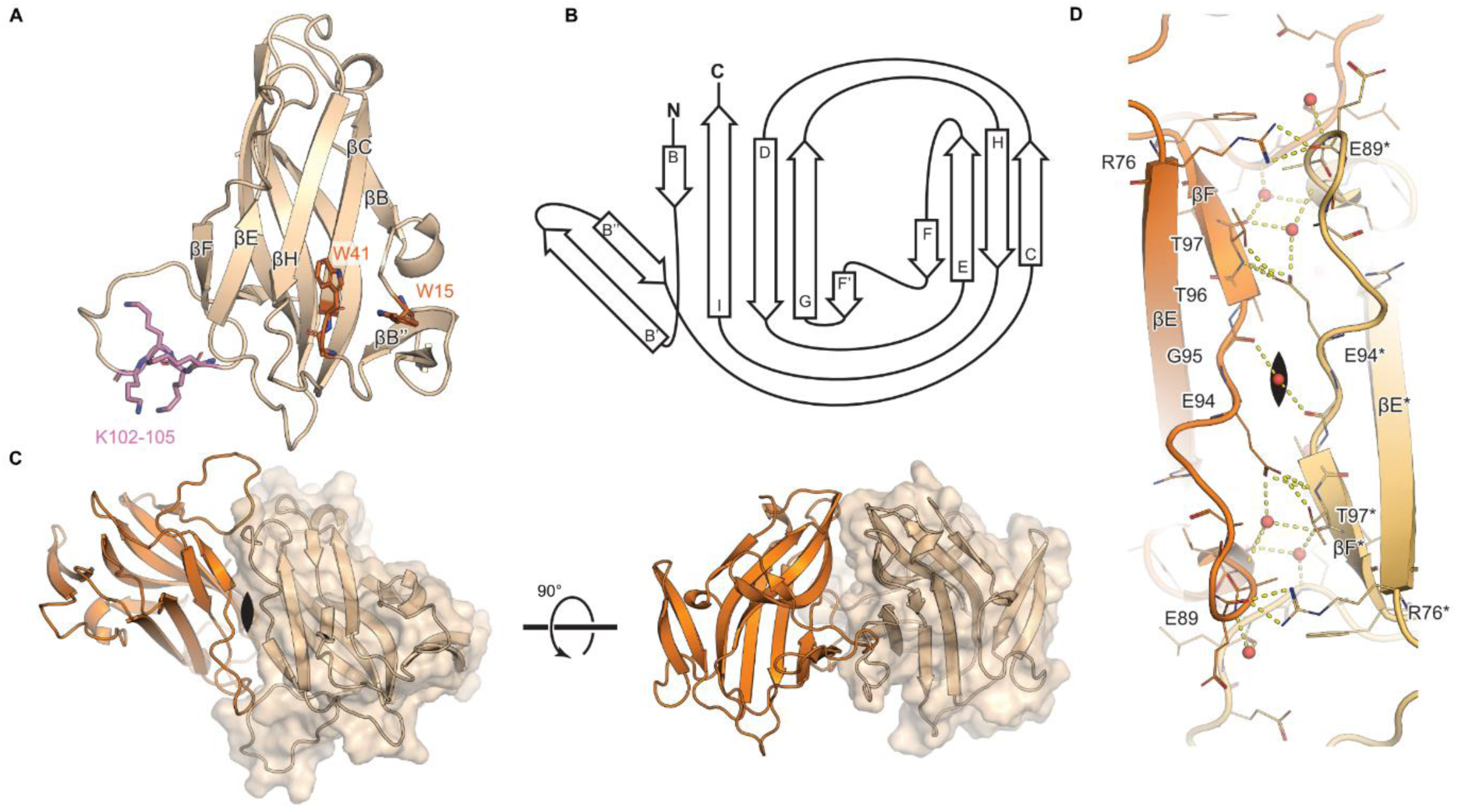
Nictaba forms a dimer of jelly rolls. A) Cartoon representation of the crystal structure of Nictaba. In pink the tetralysine motif, in orange Trp residues of special interest. B) Diagram of the Nictaba fold. C) Cartoon representation of the Nictaba dimer. One protomer is shown in orange and the second one in wheat color with a transparent surface (same orientation as in panel A). Twofold symmetry axis indicated. D) Detailed view of the dimeric interface (same orientation as in panel C). Interfacial residues shown as sticks including bridging water molecules as red spheres. Possible hydrogen-bonds are indicated by yellow dashes.

The dimerization interface mainly features interactions via hydrogen-bonding and runs along the edge of the jelly-roll fold, encompassing the E-F loops and F β-strands of the protomers (**Figure 1C,D**) and buries 940 Å² of accessible surface area which includes residues Arg76, Asn85, Glu86, Glu89, Glu94, Thr97, Gly111, Arg112, Glu128 and Phe130 (**Figure 2E**).

**Figure 2.**
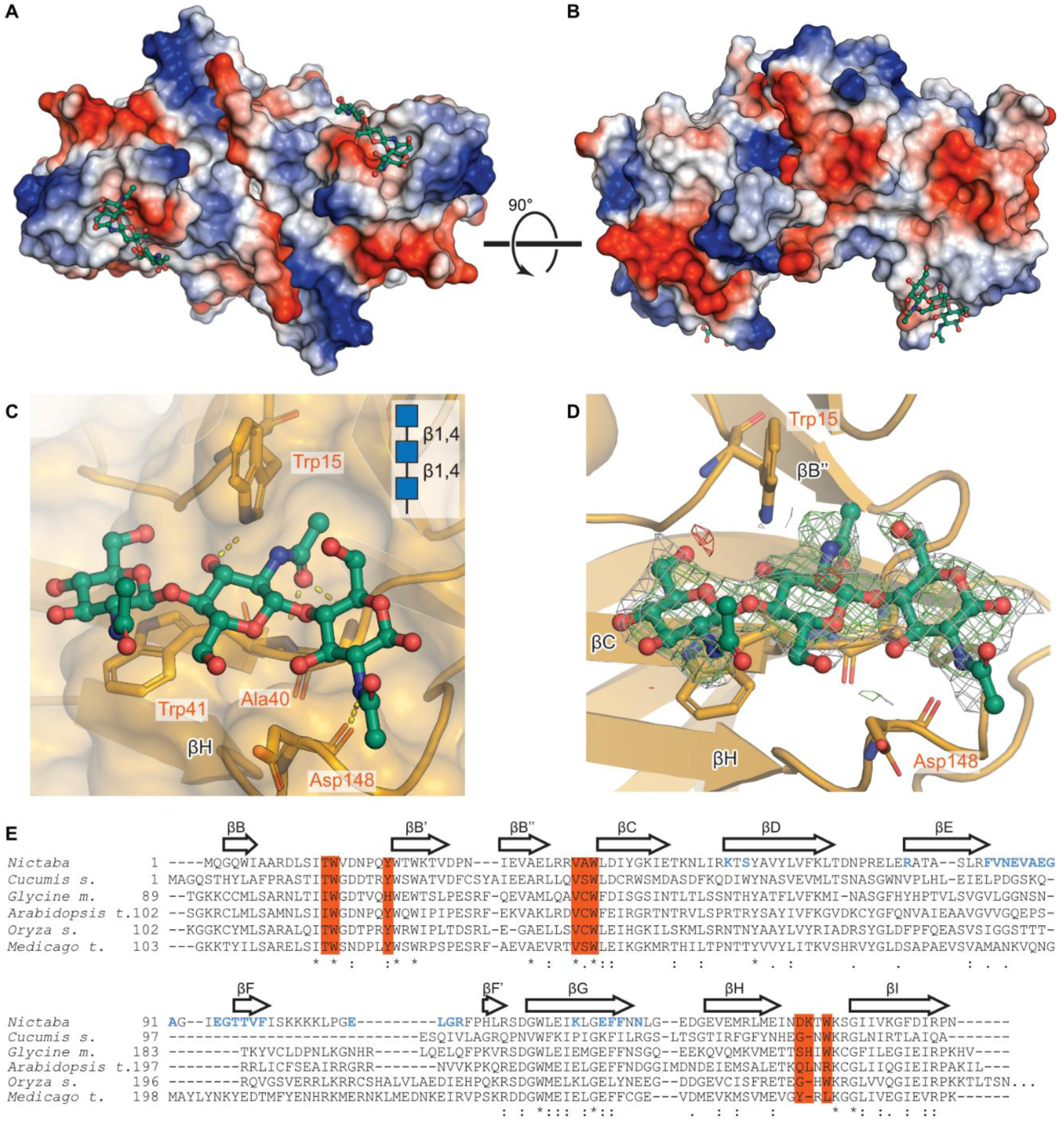
Structural basis for GlcNAc recognition by Nictaba. A-B) Surface representation of a crystal structure of a Nictaba dimer with two chitotriose molecules bound at equivalent locations. Surface coloring according to electrostatic charge, chitotriose in ball and stick representation C) Close-up view of the Nictaba chitotriose interaction. Potential H-bonds are represented as dashed yellow lines. D) Unbiased difference density carved out around the bound chitotriose moiety. In gray mesh the difference electron density with Fourier coefficients 2mFo-DFc contoured at 1σ, in green and red difference electron density Fourier coefficients mFo-DFc contoured at ± 3σ. Maps calculated by refining with torsion-angle simulated annealing in the absence of the ligands. E) Multiple sequence alignment of tobacco Nictaba and orthologs (Uniprot IDs in order of appearance Q94EW1, Q8LK69, A0A0R0GJJ1, Q9ZVQ6, B9F463 and G7I8S6). Residues of key importance to ligand binding are highlighted in orange. Residues involved in homodimerization are in blue.

The 8 copies and their dimers show little flexibility. The Residual Mean Square Deviation (RMSD) between the most and least similar dimer pairs is 0.43 Å and 0.73 Å aligning all Cα atoms and 0.24 Å and 0.56 Å when aligning all protomer Cα atoms. Some local flexibility is present at the N-termini, at the tip of the ancillary beta strand and in the tetralysine loop (Lys102-105).

Structural homology searches performed with the DALI and Foldseek servers identified CBM-35 domains (e.g. pdb 2w47, 3wnk, 3zm8) as the closest structural homologs as well as the recently published PP2 domain (pdb 7w4b). However, these do not display a similar dimerization mode. For example, the Cus17 dimer involves the N-terminal β-sheet and is further stabilized by a cysteine disulfide thereby creating a different interaction interface and dimerization mode involving the two protomers (**Supplementary Figure 3**).

### 2.2 A single GlcNAc is required for the carbohydrate-recognition of Nictaba

To obtain structural insights into the carbohydrate binding preferences of Nictaba and the structural determinants of such interactions we pursued structural studies of Nictaba in complex with chitotriose. Incubation of Nictaba crystals with chiotriose led to crystallographic electron density maps supporting the presence of chitotriose in 4 copies of Nictaba in the crystal asymmetric unit, all at the same binding site per Nictaba subunit. This binding site is partially occluded in the other 4 copies of Nictaba in the crystal asymmetric unit by crystal lattice contacts. Both protomers of one Nictaba dimer have chitotriose bound, the carbohydrate-binding sites are approximately 30 Å apart (**Figure 2A,B**).

The interaction of chitotriose to Nictaba centers around the central GlcNAc residue. Out of the 4 hydrogen bonds between the sugar and the lectin, 3 hydrogen bonds involve the central GlcNAc residue. The carbonyl oxygen of the acetyl group forms two hydrogen bonds with the backbone nitrogens of Ala40 and Trp41. These two interactions are likely responsible for the specificity of Nictaba towards GlcNAc. A third hydrogen bond is formed between the GlcNAc O3 atom and the Nε nitrogen on the Trp15 indole ring. The final hydrogen bond occurs between the first (reducing end) GlcNAc N-atom and Asp148 backbone O. The third (non-reducing end) GlcNAc residue only interacts via CH–π stacking interactions with the indole ring of Trp41 (**Figure 2C,D**). Other residues delineating the binding pocket are Thr14, Tyr21, Val39, Asp148, Lys149 and Trp151. The alignment of Nictaba with orthologs revealed conservation in residues involved in carbohydrate binding, including the key Trp15 and Trp41. Additionally, Tyr21 and Trp151as well as Val39, involved in van der Waals contacts with the ligand, are in part conserved (**Figure 2E**).

### 2.3 *In silico* simulations provide additional insights into the carbohydrate-binding in Nictaba

A combination of semi-flexible molecular docking and molecular dynamics simulations was employed to predict the binding of the Nictaba protomer with several chitooligosaccharides, an O-glycan peptide, and an N-glycan. The selected ligands (GlcNAc, GlcNAc2, GlcNAc3, GlcNAc4, GlcNAc5, O-GlcNAc peptide, and Man9GlcNAc2) were chosen based on previous knowledge about the binding capacity of Nictaba towards GlcNAc oligomers, high-mannose *N*-glycans, and O-GlcNAcylated histones. Lectin-carbohydrate systems ran for times varying from 100 ns to 250 ns, depending on the system, and the RMSD plots revealed that all systems reached equilibrium (**Supplementary Figure 4**). The intermolecular H-bonds plot over the course of the simulation can be seen in **Figure 3A**.

**Figure 3.**
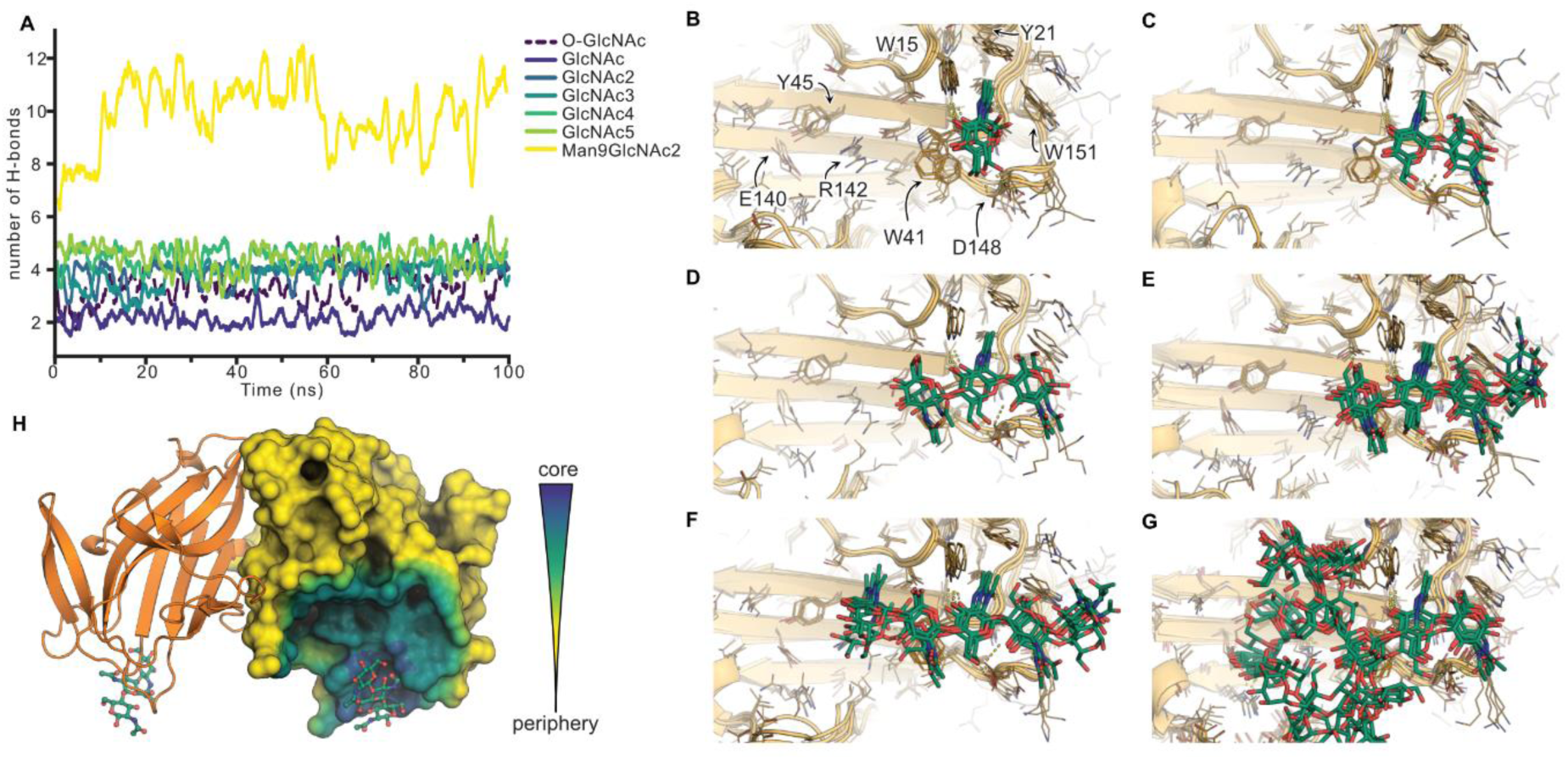
Delineation of the glycan binding site by in-silico analysis. **A)** Graph depicting the number of intermolecular hydrogen bonds formed between Nictaba and various ligands over the course of 100 ns of the molecular dynamics simulations. **B-G)** Cartoon representation of ligand poses after docking and MD. In order of appearance Nictaba in complex with: αGlcNAc, GlcNAc2, GlcNAc3, GlcNAc4, GlcNAc5 and Man9GlcNAc2. **H)** Cartoon representation depicting the extent of the glycan binding site. Same view as in Fig 2B. Surface color gradient represents if the protein is part of the core GlcNAc3 binding region (blue), likely involved in binding larger glycans at the periphery (green) or not involved in glycan recognition (yellow).

The average number of H-bonds over time increased with the size of the ligand in the following order: Man9GlcNAc2 > GlcNAc5 > GlcNAc4 > GlcNAc3 > GlcNAc2 > O-GlcNAc peptide > GlcNAc. For GlcNAc oligomers, the contacts increase significantly from GlcNAc to GlcNAc3, with GlcNAc3 to GlcNAc5 presenting marginal differences. The number of H-bonds formed between Nictaba and the Man9GlcNAc2 glycan is dramatically higher than that with any other tested ligand. For the monosaccharide, H-bonds formed between Trp15, Ala40, and Trp41 and GlcNAc seem to reach close to 100% frequency, whereas the bond between Asp148 and the O6 atom is more transient, although still more than 80% frequency (**Figure 3A**). The addition of a second GlcNAc enables the interaction of the O3 of the first GlcNAc with the backbones of Asp148 (with high frequency) and Lys149 (with 35% frequency). Additionally, weak interactions between Tyr21 and Trp151 and the first GlcNAc may play a role in the increased affinity (**Figure 3B**,**C**).

In comparison to GlcNAc2, the addition of a third monosaccharide in GlcNAc3 provides a stacking interaction with the indole ring of Trp41, as observed in the crystal structure (**Figure 3D**). The accommodation of GlcNAc4 within the binding site facilitated the formation of two extra high-frequency hydrogen bonds: one between the O6 atom of the first monosaccharide (reducing end) and Trp151, and the other between the O6 atom of the last GlcNAc (non-reducing end) and Thr41 (**Figure 3E**). For GlcNAc5, a direct H-bond between the carbonyl oxygen of the acetyl group of the last GlcNAc and Trp41 has been formed, in addition to a 15% frequency contact with Arg142 (**Figure 3F**).

The interaction with the Man9GlcNAc2 glycan involves several additional interactions in comparison to the tested GlcNAc oligomers and has been applied in the current work to identify the extended binding site of the lectin. The glycan is placed on the binding site through its chitobiose core. The Manβ1,4-GlcNAcβ1,4-GlcNAc substructure fits within the binding site in a similar position as GlcNAc3 with the beta-mannosyl kept in place through a stacking interaction with Trp41. Besides the recurrent interactions observed for GlcNAc3, the following high-frequency contacts were also observed: Ser12, Tyr45, Arg82, Gly90, Ala91, Glu140 (**Figure 3G**).

The interactions with the O-GlcNAc peptide follow different principles compared to the binding of GlcNAc. Trp41 forms a stacking interaction and a high-frequency H-bond with the O3 atom of GlcNAc, whereas Trp15 also interacts with the O3 atom and the acetyl O atom. The peptide portion of the glycan interacts with high frequency with Asp148, and low (11%) frequency with Lys149 (**Supplementary Figure 5**).

### 2.4 Glycan array and structural data provide additional insights into Nictaba carbohydrate binding

Nictaba at concentrations of 50 µg/mL or 200 µg/mL was previously tested on the version 5.0 of CFG glycan array yielding consistent results (Blixt et al., 2004; Smith et al., 2010). For chito-oligosaccharides, the recognition hierarchy is as follows: GlcNAc5 > GlcNAc6 > GlcNAc3 (glycans 190-192). Apart from chito-oligosaccharides, Nictaba exhibited strong binding to the Man3GlcNAc2 glycan (glycans 50 and 51), which forms the core of *N*-glycans. Motif analysis revealed two primary motifs: the core Man2GlcNAc2, with a relative binding of 72%, and GlcNAcβ1-4GlcNAcβ1-4GlcNAc with a relative binding of 100%. A third motif, Galb1-4GlcNAcβ1-2Manα1-6Manβ1-4GlcNAcβ1-4GlcNAc, found in some complex glycans, exhibited lower relative binding at 39% (**Table 2**).

**Table 2.**
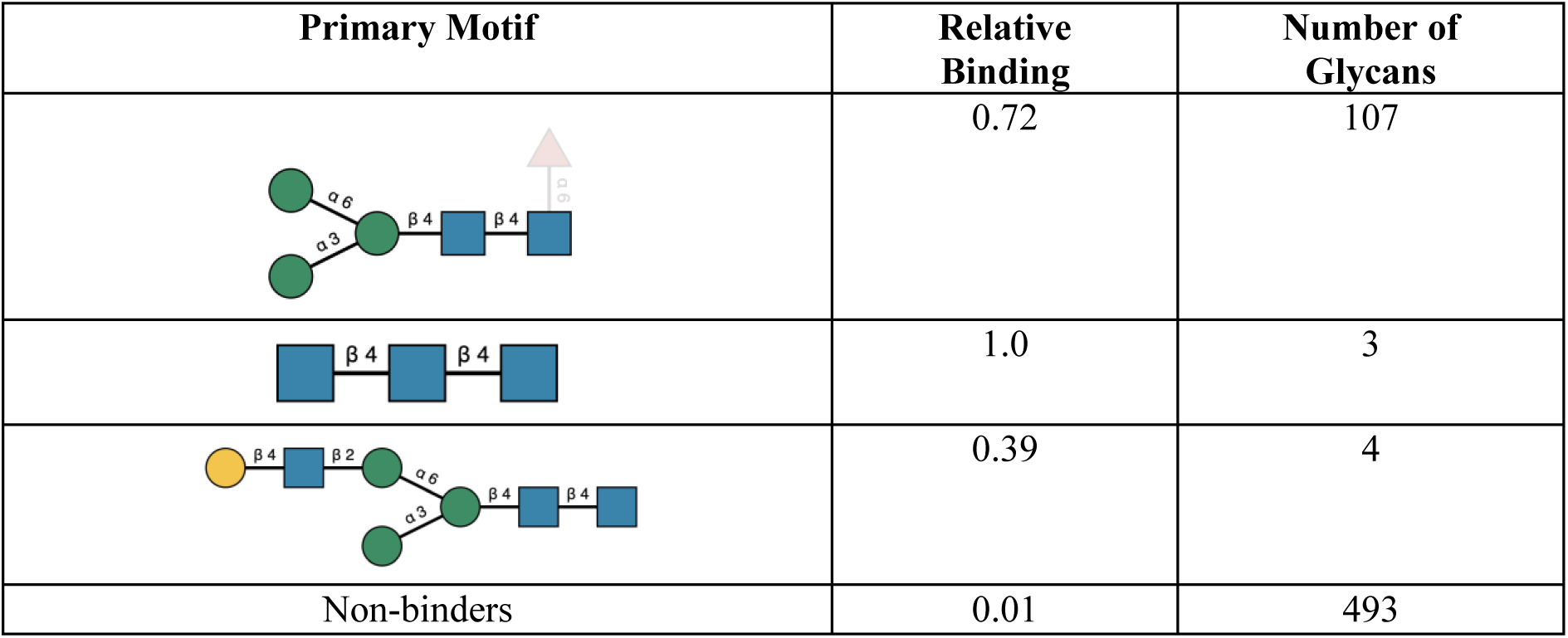
Glycan binding motifs identified for Nictaba by CarboGroove within the CFG array v5.0 using the dataset of Nictaba at 200 µg/mL. Sugars are depicted using the Symbol Nomenclature for Glycans (SNFG).

For N-glycans, MCAW-DB results indicate that Nictaba exhibits limited tolerance to modifications within the Man3GlcNAc2 core for binding (**Supplementary Figure 6**). Fucα1-6GlcNAc is the only one modification that is well-tolerated. Docking analysis suggests that this fucosylation does not alter the ligand’s position within the binding site and can form an additional H-bond with Tyr21, potentially enhancing binding. This is supported by the strongest binder having this substructure (glycan 473). Fucα1-3GlcNAc is not beneficial due to poor contact with Lys149. Fucosylations on the second GlcNAc of the core result in clashes with Tyr21, Val39, Ala40, Trp41, and Trp151, rendering the interaction impossible. Sialylation is generally unfavorable for binding, although α2-6 sialylation within the branches starting with the 1,6 mannosyl arm of Man3GlcNAc2 is better tolerated than that of the α1-3 mannosyl arm.

## 3. DISCUSSION

Nictaba is a stress-inducible lectin whose expression is upregulated in *Nicotiana tabacum* leaves and roots after exposure to some stress conditions (Eggermont et al., 2017). Previous data revealed that Nictaba is a nucleocytoplasmic lectin specific for GlcNAc oligomers and capable of interacting with several *N*-glycans as well as O-GlcNAc modified histone proteins (Delporte et al., 2014; Lannoo et al., 2006). Despite being studied for a long time, the experimental structure of Nictaba remained unknown. To address this gap, the current work delved into the crystal structure of this lectin.

The Nictaba monomer is formed by two antiparallel β-sheets parallel to each other forming a jelly-roll fold. Each sheet is formed by four β-strands interlinked by loops of variable sizes, with a particularly large, exposed loop containing the tetralysine motif known to be important for the nuclear localization of the lectin (Lannoo et al., 2006). A β-hairpin substructure can also be found near one of the sheets. The observed jelly-roll fold is not uncommon among plant lectins, with several families, including the *Agaricus bisporus* agglutinin family and the well-researched legume lectin family, presenting the same overall fold (Ismaya et al., 2020; Loris et al., 1998). However, the specific architecture of Nictaba and its mode of carbohydrate recognition are uncommon. A similar crystal structure was reported recently for the phloem lectin from *Cucumis sativus* (Cus17) (PDB id: 7w4b) (Bobbili et al., 2023). Monomer superposition results in an RMSD of 1.06 Å, a value that correlates with a very high degree of structural similarity despite the 24% sequence identity. For the remaining PDB structures, a search through PDBeFold (Krissinel & Henrick, 2005) resulted in few results with most being variations of the CBM35 proteins from *Clostridium thermocellum* (PDB id: 2w47) displaying RMSDs of 2.93 Å.

The Nictaba dimer is held together by several H-bonds formed between the residues Arg76, Glu85, Glu89, Glu94, Thr96, Thr97, Gly111, Arg112, Glu128, and Phe130, present on the edge of the loops E-F and F β-strands of the protomers (**Figure 1D**). When comparing the Nictaba dimer with other structures within the PDB, no matches have been found, thus revealing a new dimerization interface within plant lectins.

Specific carbohydrates interact with and are bound to the carbohydrate-recognition domain of a lectin through a combination of H-bonds, hydrophobic, van der Waals and other weak interactions (Bojar et al., 2022; Zhang et al., 2021). Our crystallographic data revealed that Nictaba possesses a single binding site per monomer. The carbohydrate-recognition domain forms a shallow cavity on the lectin surface, composed of three loops. The primary binding site comprises the following residues: Thr14, Trp15, Tyr21, Val39, Ala40, Trp41, Asp148, Lys149, and Trp151. The interaction network mainly centers around the central GlcNAc molecule of chitotriose and is mediated by a maximum of four H-bonds and additional hydrophobic, stacking and van der Waals interactions. Specifically, the residues Ala40, Trp41, and Asp148 directly interact with the O- and N-atoms of the acetamide groups and can be considered the key determinants for the specificity of Nictaba (**Figure 2C,D**). In contrast to other plant lectins, no water bridges were observed for the carbohydrate interaction. The binding profile to GlcNAc3 is quite similar to Cus17, with some differences that can be attributed to map interpretation. However, Nictaba features interactions at the O6 position that are absent in Cus17. Additionally, in Cus17 and other similar phloem proteins from *Cucurbitaceae*, a serine residue replaces the Ala40 residue found in Nictaba, which is unlikely to make a major difference. When comparing the amino acid sequence of Nictaba to other similar lectins, including *Cucumis sativus* PP2 protein (UNIPROT id: A0A0A0KYP0); TIR-domain containing protein from *Arabidopsis thaliana* (UNIPROT id: Q8H7D6); *Cucurbita maxima* PP2 protein (UNIPROT id: Q42383); F-box family protein from *Ricinus communis* (UNIPROT id: B9RFT4); F-box containing protein from *Oryza sativa* (UNIPROT id: Q6K3F5) and phloem lectin from *Zea mays* UNIPROT id: B6TVP3), the residues Trp15, Tyr21, Val39, Trp41, and Trp151, all part of the binding site, appear to be conserved.

The interaction between Nictaba and GlcNAc, as well as chitooligosaccharides (GlcNAc2, GlcNAc3, GlcNAc4, and GlcNAc5), has been investigated in more detail through *in silico* simulations. Molecular dynamics (MD) simulations revealed stable interactions with all tested ligands (**Figure 3A**). Specific interactions have been identified and are detailed in section 2.3. Previous studies have already demonstrated an increased affinity of Nictaba for longer chitooligosaccharides. It has been shown that the affinity of the lectin towards GlcNAc2, GlcNAc3, and GlcNAc4 is 50, 1600, and 3000 times higher compared to its affinity for GlcNAc, respectively (Chen et al., 2002). This enhanced affinity can be attributed to the additional interactions and/or increased contact frequency with some residues resulting from the increased ligand size. Interestingly, due to the shape of the binding site and the linear structure of chitooligosaccharides, the differences in interactions between GlcNAc3, GlcNAc4, and GlcNAc5 are small. This phenomenon has also been observed in Surface Plasmon Resonance experiments, where GlcNAc3 and GlcNAc4 exhibited close binding affinities. Hence, it is generally accepted that GlcNAc3 is the most complementary ligand for Nictaba’s binding site, this being in line with other related lectins (Beneteau et al., 2010; Mondal & Swamy, 2020).

As the interaction mode between Nictaba and its ligand had not yet been observed when we solved the crystal structure, we submitted it to the Critical Assessment of Techniques for Protein Structure Prediction 15 (CASP15) (Robin et al., 2023) competition as part of the ligand binding prediction (T1187). All entries failed to identify a binding pose resembling the experimental observation we provide in this paper (Best Local Distance Difference Test for Protein–Ligand Interactions score 0.385). This failure to identify the ligand binding site by *in silico* methods stands in stark contrast to the very similar experimentally observed ligand pose for the Cus17 chitotriose cocrystal structure and underlines the value of experimental structural biology.

Glycan array data already highlighted the interaction of Nictaba with several high-mannose and complex *N*-glycans (Schouppe et al., 2010). In this study, we leveraged this data in conjunction with protein bioinformatics to identify the structural aspects of Nictaba binding to glycans. This analysis allowed us to pinpoint the primary binding motifs within the array, which are the chitobiose core of *N*-glycans and, unsurprisingly, GlcNAcβ1-4GlcNAcβ1-4GlcNAc. This specificity profile towards *N*-glycans aligns with that of other Nictaba-related lectins, including PP2 lectins from *Arabidopsis thaliana*, *Cururbitaceae* species, etc. (Allen, 1979; Anantharam et al., 1986; Beneteau et al., 2010; Narahari et al., 2011). Already in 1979 Allen showed the interaction of the *Cucurbita pepo* lectin with glycopeptides (derived from fetuin and soybean agglutinin) carrying high mannose and complex *N*-glycans (Allen, 1979). Anantharam et al. (1986) provided a detailed analysis of the chito-oligosaccharide specific lectin from the exudate of ridge gourd *(Luffa acutangula)* fruits and showed that the chitobiosyl core of *N*-glycans is indispensable for their interaction with the lectin since EndoH-treated glycopeptides of soybean agglutinin do not bind to the *Luffa* lectin (Anantharam et al., 1986). Molecular docking and MD analysis, using the high-mannose glycan Man9GlcNAc2 as a ligand, provided insights into the potential binding mode of Nictaba to *N*-glycans. The binding was found to be highly stable, with the Man3GlcNAc2 core occupying the primary binding site and the beta-mannosyl being stabilized through a stacking interaction with Trp41. The remaining mannosides were stabilized by the extended binding site of Nictaba (**Figure 3H**). The presence of an extended binding site is a common property of plant lectins and is seen in different families (Asensio et al., 2000; Cavada et al., 2022). Furthermore, glycan array data revealed that Nictaba exhibits a 2-fold higher affinity towards the core Man3GlcNAc2 compared to GlcNAc3. Additionally, molecular docking analysis showed no difference in the interaction scores between GlcNAc3 and Manβ1-4GlcNAcβ1-4GlcNAc. This similarity arises from the fact that the mannosyl moiety is stabilized by CH–π stacking interactions with the indole ring of Trp41, an interaction independent of the acetyl group of GlcNAc.

Studies applying different approaches attempted to investigate the interaction of Nictaba with nuclear histones. The first study utilized Nictaba-immobilized affinity chromatography to capture nuclear proteins from Tobacco cv Xanthi cells. The lectin successfully captured the core histone proteins H2A, H2B, H3, and H4. Bound histones were released from the Nictaba column upon elution with 1M GlcNAc, suggesting a lectin-glycan interaction (Schouppe et al., 2011). In the second study, bimolecular fluorescence complementation (BiFC) assays confirmed the interaction between Nictaba and H2A, H2B and H4 histones from *Arabidopsis thaliana* (Delporte et al., 2014). Given the hypothesis that the lectin interacts with histone proteins through an O-GlcNAc modification (proven by western blotting using antibody directed against O-GlcNAc modification on Ser/Thr residues), we conducted *in silico* binding studies to evaluate the binding mode and compare it to known ligands. The interaction between Nictaba and O-GlcNAc was found to be stable and primarily utilized the same residues in the binding site as for GlcNAc. However, the sugar portion of the glycan shifted its position to engage in stacking interactions with Trp41 and form hydrogen bonds with Trp41 and Trp15, while the peptide portion established interactions with Asp148, and, to a lesser extent, Lys149.

The interaction of Nictaba with O-GlcNAcylated glycoproteins suggests its involvement in signal transduction regulation and/or modulation, potentially altering responses related to stress in plants (Xu et al., 2017). This observation aligns with the presence of Nictaba-like proteins in the transcriptomes of various plants, including *Arabidopsis thaliana*, tomato, cucumber, and rice, which have been associated with plant defense and stress responses (Van Holle et al., 2017; Wojszko et al., 2023).

In conclusion, the current study provided a comprehensive structural analysis of the stress-inducible lectin from *Nicotiana tabacum*, detailing its structural, biochemical, and functional properties through experimental and computational methods. The discovery of Nictaba’s structure, unique oligomerization mode, carbohydrate-binding specificity, and interaction with various glycans can enhance our understanding of the physiological role of the lectin in stress responses but can also establish a foundation for further exploration of lectin-mediated cellular processes and the potential of Nictaba for biotechnological applications.

## 4. MATERIALS AND METHODS

### 4.1 Constructs

DNA encoding Nictaba (Uniprot Q94EW1) was codon optimized for expression in *E.coli* and synthesized by Twist Bioscience (San Francisco, CA, USA). To allow easy cloning into the pET28 vector the 5’ end was modified to encode an NcoI site. An N-terminal His-tagged version of the construct was cloned by PCR. A hexahistidine tag followed by a caspase 3 recognition site (DEVD) was added in frame to the 5’ end and subcloned into pET28.

### 4.2 Expression and Purification

The pET28 vector, encoding Nictaba, was introduced into chemically competent *E. coli* strain BL21 DE3 pLysIq (NEB no. C3013) through heat shock. Transformed colonies were selected on Luria Broth agar plates (2% w/v agar) containing kanamycin (25 µg/ml). An overnight preculture, cultivated in Luria Broth medium with kanamycin (25 µg/ml), was used to inoculate the expression culture in Luria Broth medium supplemented with 25 mM phosphate at pH 7.4 and kanamycin (5% v/v). The expression culture was cultivated at 36 °C until reaching an OD600nm of approximately 0.6. At this point, induction was initiated by adding 0.5 mM IPTG, and the temperature was reduced to 20 °C for overnight expression. Cells were harvested through centrifugation, and the pellet was lysed by two passes through a high-pressure homogenizer in the presence of DNase. The supernatant fraction was then clarified through centrifugation at 25,000 g, followed by filtration.

Untagged Nictaba was purified in five steps as outlined below: Firstly, the clarified supernatant was supplemented with 25 mM PIPES, and the pH was adjusted to 6.0, with conductivity set to <10 mS/cm. Subsequently, cation exchange chromatography (C-IEX) was conducted on a SP Sepharose resin at pH 6.0, resulting in the detection of Nictaba in the flow-through fraction. The lectin-containing fraction was then enriched with 25 mM acetate, and the pH was further adjusted to 5.0, causing notable precipitation. An additional clarification step involving centrifugation and filtration was necessary before a second cation exchange separation on SP Sepharose. Nictaba bound to the resin and eluted with approximately 150 mM NaCl in the elution buffer. The eluate containing Nictaba was neutralized by the addition of HEPES buffer and further purified through Size-Exclusion-Chromatography (SEC) on an SD200 16/600 column equilibrated with HBS (25 mM HEPES pH 7.4, 150 mM NaCl). A peak at 84 ml retention volume contained the lectin. To enhance purity, anion exchange chromatography (A-IEX) on Mono Q resin was attempted. The sample was supplemented with 25 mM Tris at pH 8.0, and conductivity was adjusted to <10 mS/cm. Under these conditions, Nictaba did not bind to the resin and remained in the flow-through (FT). SEC was employed as a final polishing step, where the A-IEX flow-through was concentrated and loaded onto an SD200 Increase 10/300 column equilibrated with HBS. Nictaba eluted as a single peak at a retention volume of 15 ml.

Purification of His-tagged Nictaba involved immobilized metal affinity chromatography (IMAC) followed by SEC. The His-tag was removed through overnight incubation of the protein with recombinant Caspase3 enzyme (1% w/w). Tagged, undigested protein was separated from the cleaved protein by IMAC. As a final polishing step, the IMAC flow-through was concentrated and loaded onto an SD200 Increase 10/300 column equilibrated with HBS.

### 4.3 Crystallization and Soaking

Extensive screening was conducted using commercial sparse matrix screens from Hampton Research (Aliso Viejo, CA, USA) and Molecular Dimensions (Sheffield, UK) in the sitting drop vapor diffusion geometry. Two crystalline hits were identified under the Proplex F8 (1.0 M (NH_4_)_2_SO_4_, 100 mM NaAc pH 5.0) and BCS screen E3 (100 mM ZnAc, 100 mM ZnCl2, 100 mM Bis-Tris pH 7.5, 20% Medium smear) conditions. The presence of an 18 kDa protein in the crystals from both conditions was confirmed by loading the crystals onto SDS-PAGE. Optimization efforts were successful only for the BCS E3 condition, resulting in well-diffracting crystals.

For the Nictaba crystal structure in complex with chitotriose, crystallization drops containing crystals were supplemented with mother liquor enriched with 5% (v/v) glycerol, 5% (v/v) ethylene glycol, and 5% (v/v) DMSO. The DMSO included a stock of chitotriose at 20 mM, yielding a 1 mM chitotriose solution. Crystals were allowed to soak for approximately 45 minutes before vitrification.

### 4.4 X-ray Diffraction Data Collection and Structure Solving

A single crystal of apo-Nictaba, grown in 91 mM ZnAc, 91 mM ZnCl_2_, 91 mM Bis-Tris pH 7.5, and 18.2% Medium PEG smear, was cryoprotected with the addition of 17.5% (v/v) ZW221 to the mother liquor before being vitrified in liquid nitrogen. The diffraction data were collected at the EMBL P14 beamline (Petra3, Hamburg, DE) at 100 K, and the crystal diffracted to a resolution of 2.15 Å. The data were integrated and scaled using XDS, belonging to space group P 2_1_ with unit cell dimensions a=66.51 Å, b=145.97 Å, c=82.05 Å, and β=95.95°. Full dataset statistics are reported in Table 1. Similar to the apo-Nictaba crystal, the GlcNAc3-soaked Nictaba crystal was grown from the same mother liquor. After vitrification, X-ray diffraction data were measured at the EMBL P14 beamline, and the crystal diffracted to 2.0 Å, showing isomorphism.

Molecular replacement in Phaser, as implemented in Phenix, utilizing assorted models derived from proteins proposed as structural homologues (pdb depositions 1cx1, 1gu3, 2y6h, 3k4z, 3p6q, 4mgq), yielded TFZ scores between 5.5 and 6.9, none of which were interpretable.

Phases were recovered by Single-wavelength Anomalous Diffraction (SAD). Since phases could not be readily recovered from the anomalous signal of Zn atoms in the crystallization condition, iodine soaks were performed. A single crystal of approximately 50 x 50 x 10 µm was transferred from mother liquor to a drop of 50 mM ZnAc, 50 mM ZnCl2, 60 mM BisTris, 40 mM Tris pH 7.5, 400 mM NaI, 20% PEG Smear Medium, and 15% ZW221 cryo. After approximately 2 minutes, it was scooped into a Mitigen dual thickness Microloop LD loop and vitrified in liquid nitrogen.

Five datasets with 360° rotation were collected at 6 keV energy at full flux and 0.15° oscillation per frame. The kappa angle of the goniometer was altered between the 5 dataset collections to minimize instrumental errors. The data were integrated and scaled using XDS and XSCALE, with an overall anomalous multiplicity of 16.87. The crystal belongs to the same space group but does not appear to be isomorphous (Unit cell dimensions a=69.47 Å, b=145.71 Å, c=82.08 Å, β=96.75°) compared to our best-diffracting apo crystal.

Substructure solution and initial model building were carried out using the CRANK2 pipeline, as implemented in the CCP4 online web service (Krissinel et al., 2018; Skubák & Pannu, 2013). The substructure solution comprised 4 atoms with near full occupancy (actual iodine atoms) and over 20 atoms with half and lower occupancy (potential zinc atoms). The CRANK2 pipeline constructed 8 nearly complete and accurate Nictaba molecules arranged as 4 dimers. One such dimer was employed to phase the higher resolution non-isomorphous dataset through maximum-likelihood molecular replacement in Phaser, as implemented in the Phenix suite (Translation function Z-score (TFZ) = 91.0, Log-Likelihood-Gain (LLG) =11246).

Similarly, phases could be retrieved by the CRANK2 pipeline from a europium-soaked crystal. For this, a single crystal was soaked in mother liquor supplemented with 100 mM Eu coordinated by 1,4,7,10-tetraazacyclododecan-1,4,7-triacetic acid (DO3A) and 17.5% ZW221. After 2 minutes of soaking, the crystal was rinsed in mother liquor supplemented with 17.5% ZW221 and vitrified in liquid nitrogen. A redundant dataset was collected at 8.09 keV (Table 1). Similarly to the previous monoclinic crystals, this one was also non-isomorphous, with unit cell constants of a=70.16 Å, b=151.42 Å, c= 82.51 Å, and β= 97.63°. While still readily interpretable, the model output by the CRANK2 pipeline was of lower quality compared to the one obtained from Iodine derivatization.

Iterative model building and refinement were conducted in Coot (Casañal et al., 2020) and Phenix.refine (Liebschner et al., 2019). The models and datasets were deposited in the Protein Data Bank under accession numbers 8qmg and 8ad2for apo and NAG3 soaked proteins, respectively. The integrated diffraction data of the iodine and europeum soaks are accesible as part of the deposited apo dataset. Model statistics are reported in Table 1. Structural analysis was performed in PyMol (Schrodinger LLC, USA). and UCSF Chimera (Pettersen et al., 2004).

### 4.5 Molecular Docking and Molecular Dynamics Simulations

The interaction between Nictaba and carbohydrate structures (GlcNAc, diacetylchitobiose (GlcNAc2), triacetylchitotriose (GlcNAc3), tetraacetyl-chitotetraose (GlcNAc4), pentaacetyl-chitopentaose (GlcNAc5), the Man9GlcNAc2 glycan, and an O-GlcNAc peptide) has been predicted through a combination of semi-flexible molecular docking and explicit solvent molecular dynamics (MD). Inputs for molecular docking included the structure file of Nictaba in complex with GlcNAc3 and the respective ligand files. Ligands were designed using the carbohydrate-builder module of Glycam-Web (http://glycam.org), except for the O-GlcNAc peptide, which was designed in the glycoprotein-builder of the same server; the GlcNAc was attached to the Ser of the PPVSEASST sequence.

The complex formation was predicted in GOLD v.2023 (Jones et al., 1997) with default settings and ChemPLP as scoring function (Korb et al., 2009). Receptor and ligands were prepared by removing solvent molecules and adding missing hydrogen atoms. Ligand fitting was restricted to the binding site, and all residues within an 11 Å radius. The receptor was kept rigid whereas the ligand was made flexible. The best pose for each ligand was selected considering a combination of docking score, expected H-bonds, and ligand geometry distortions. Redocking with GlcNAc3 has been used to assess the algorithm suitability for the current application.

The best docking poses were utilized as inputs for MD simulations. The systems were prepared in CHARMM-GUI using the parameters of the CHARMM36m force field (Huang et al., 2017). All systems were solvated using the TIP3P water model and neutralized with Na+ counter ions. Simulations were conducted in the pmemd.cuda module of the AMBER22 suite (Case et al., 2022) following a pipeline that involved energy minimization through a steepest descent algorithm with 10 kJ/mol as the convergence criterion. Equilibration in 250,000 steps of the NVT ensemble, and subsequently, the same number of steps under the NPT ensemble. After equilibration, the MD was set to run under the NPT ensemble for times varying depending on the time required for the systems to fully transition to the solvated state, with a minimum of 100 ns under a time step of 2 fs. The SHAKE algorithm (Ryckaert et al., 1977) was applied to constrain covalent bonds involving hydrogen atoms. The Particle Mesh Ewald (PME) method (Essmann et al., 1995) was employed to calculate long-distance electrostatic interactions with a threshold of 9 Å. Temperature and pressure parameters were set to 300 K and 1 bar, respectively, kept stable by a Langevin thermostat and an isotropic Monte-Carlo barostat (Berendsen et al., 1984).

Trajectory equilibrium was accessed through protein backbone root-mean-square deviation (RMSD) values. Lectin-carbohydrate interaction properties were studied by a combination of intermolecular H-bonds and contact frequencies. The Cpptraj module of AmberTools23 was used for the analysis across all systems (Roe & Cheatham, 2013), Xmgrace, and VMD have been applied for graph preparation and trajectory visualization respectively (Humphrey et al., 1996).

### 4.6 Glycan array analysis

Glycan array data for native Nictaba purified from jasmonate-treated leaves (Chen et al., 2002) or recombinant lectin (Schouppe et al., 2010) are publicly available within the Consortium of Functional Glycomics data section (http://www.functionalglycomics.org). For the analysis, the glycan array motif analysis of CarboGroove (Klamer et al., 2022) and MCAW-DB (Hosoda et al., 2018) has been used in combination with molecular docking, as explained in section 4.5, and manual analysis.

## ACKNOWLEDGEMENTS

The synchrotron data was collected at beamline P14 operated by EMBL Hamburg at the PETRA III storage ring (DESY, Hamburg, Germany). We would like to thank Isabel Bento for the assistance in using the beamline. We would like to thank José Martins for the gift of chitotriose used in structure determination. The authors thank the Consortium of Functional Glycomics (CFG) for the help with the glycan array data. S.N.S. acknowledges research support from the Flanders Institute for Biotechnology (VIB grant C0101). Y.B. was a post-doctoral research fellow supported by the Research Foundation Flanders (FWO grant no. 12S0519N) and is an ARISE fellow funded from the European Union’s Horizon 2020 research and innovation programme under the Marie Skłodowska-Curie grant agreement No 945405. V.J.S.O is a postdoctoral research fellow supported by the Research Foundation Flanders (FWO grant no. 12T4622N).

## AUTHOR CONTRIBUTION STATEMENTS

Yehudi Bloch (Conceptualization, Methodology, Investigation, Visualization, Writing – original draft, Writing – review & editing). Vinicius Jose Da Silva Osterne (Conceptualization, Investigation, Visualization, Writing – original draft, Writing – review & editing). Savvas N. Savvides (Conceptualization, Validation, Writing – original draft, Writing – review & editing, Supervision and Funding acquisition). Els J.M. Van Damme (Conceptualization, Validation, Writing – original draft, Writing – review & editing, Supervision and Funding acquisition).

## DATA AVAILABILITY STATEMENT

The data underlying this article will be shared on reasonable request to the corresponding author.

