## Supplementary Figure 1 for "The crystal structure of Nictaba reveals its carbohydrate-binding properties and a new lectin dimerization mode"

**Supplementary Figure 1.** SEC-MALLS and SDS-PAGE analysis of the purified Nictaba. A) SEC-MALLS chromatogram (orange) and MW determination (blue) of Nictaba same as in subpanel B lane 5. B) SDS-PAGE analysis of the purified tagless and His-tagged Nictaba. Lane 1, final purified material of tagless Nictaba as utilized in crystallization trials. Lanes 2,3,4,5 His-tagged Nictaba after IMAC, after caspase digest, flow-through from IMAC and final purified material of cleaved Nictaba.

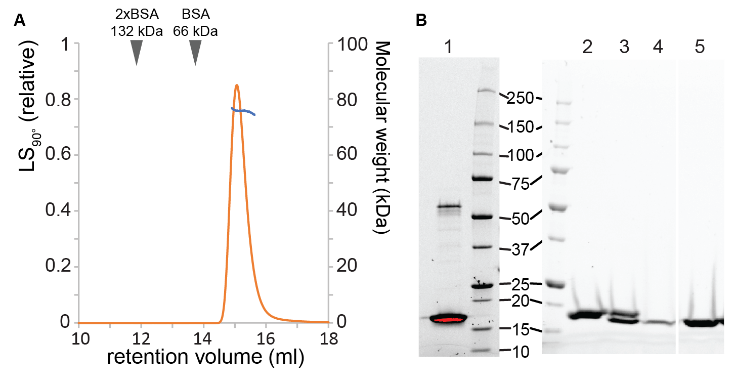
