## Supplementary Figure 2 for "The crystal structure of Nictaba reveals its carbohydrate-binding properties and a new lectin dimerization mode"

**Supplementary Figure 2.** Anomalous signal is present in the SAD datasets of both Iodine and Eu-DO3A soaks. A-C) Anomalous signal to noise, cross-correlation and map coming from the NaI soaked crystal C) Anomalous electron density map carved 5 Å around one Nictaba dimer contoured at 12σ. D-F) Anomalous signal to noise, cross-correlation and map coming from the Eu-DO3A soaked crystal F) Anomalous electron density map carved 5 Å around one Nictaba dimer contoured at 8σ.

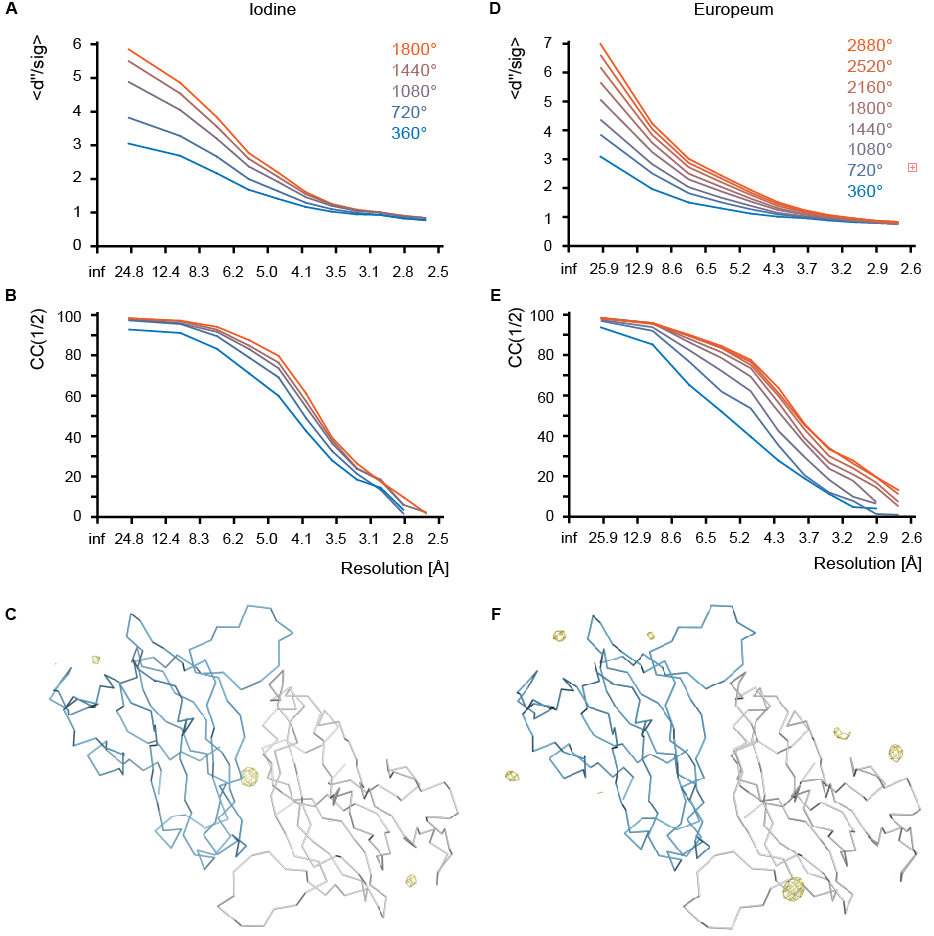
