## Supplementary Figure 3 for "The crystal structure of Nictaba reveals its carbohydrate-binding properties and a new lectin dimerization mode"

**Supplementary Figure 3.** Nictaba and Cus17 form distinct dimers. A) Cartoon representation of the structural alignment of Nictaba and Cus17 dimers showing the opposite orientation of the second protomer. B) Detailed view of the Cus17 dimerization interface centered around the N-terminus. Potential hydrogen bonds indicated as yellow dashed lines. C) Multiple sequence alignment of Nictaba and Cus17 related proteins by ClustalOmega. In order of appearance Uniport sequences Q94EW1,P0DSP5,A0A1S3CE66,Q8L4J1. Marked in blue box, the Cus17 N-terminal region and with a yellow box, the Cys residue which form the Cus17 dimerization interface. Marked with orange boxes, the residues involved in ligand binding. In blue font the residues defining the Nictaba dimerization interface.

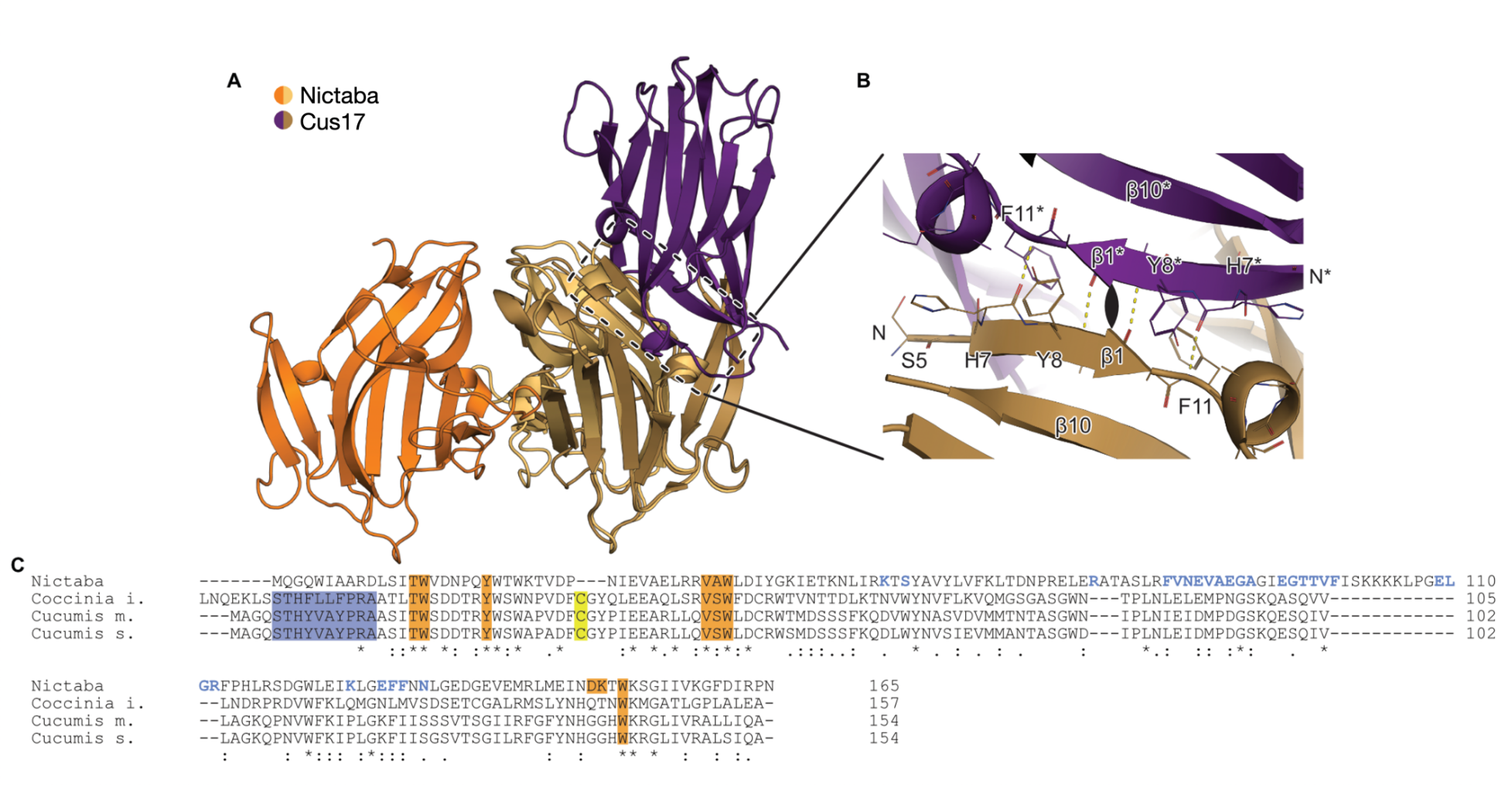
