## Supplementary Figure 4 for "The crystal structure of Nictaba reveals its carbohydrate-binding properties and a new lectin dimerization mode"

**Supplementary Figure 4.** Root-mean square deviation of the protein atoms over the course of the simulations.

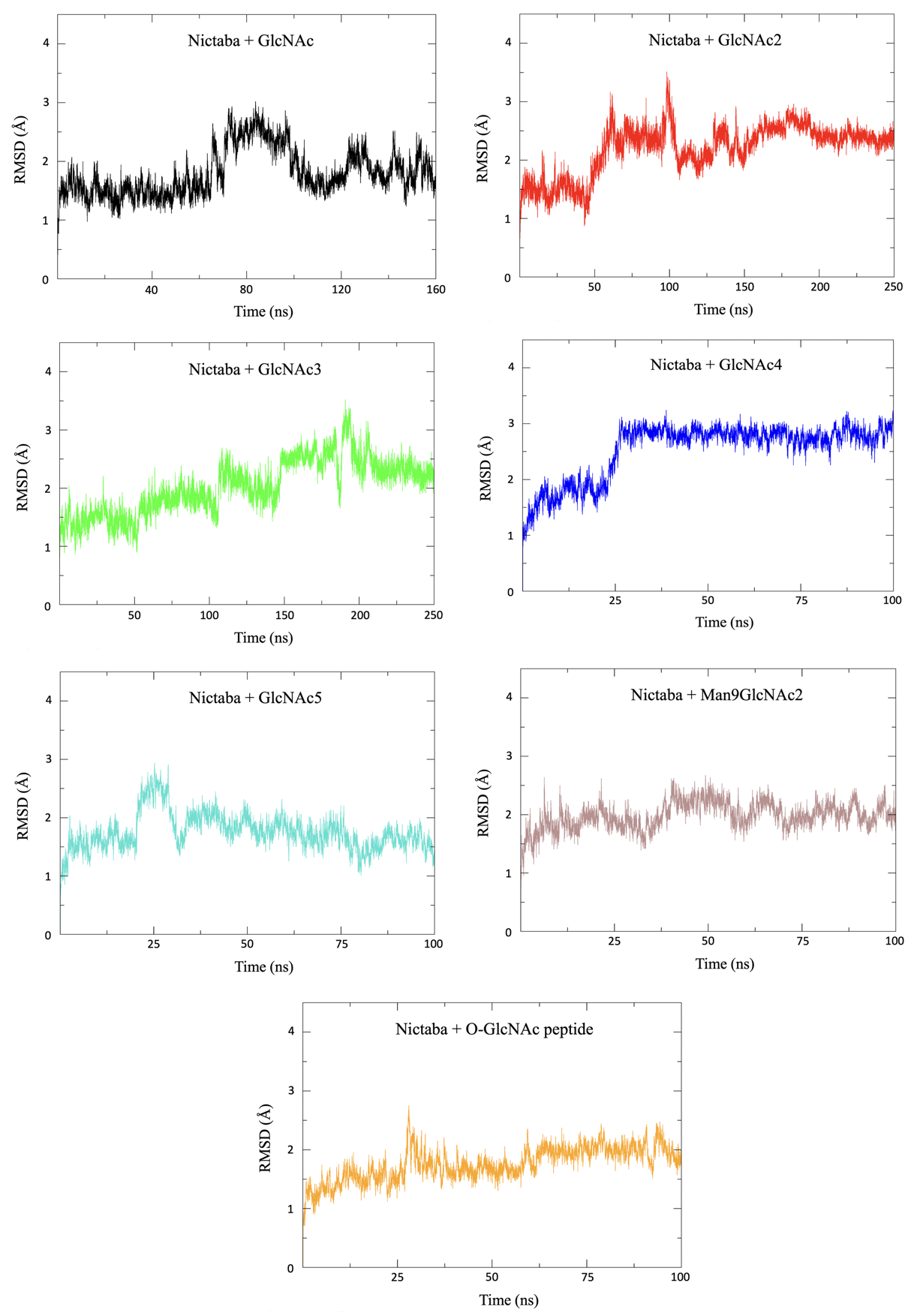
