## Supplementary Figure 5 for "The crystal structure of Nictaba reveals its carbohydrate-binding properties and a new lectin dimerization mode"

**Supplementary Figure 5.** Interactions formed between Nictaba and the O-GlcNAc peptide. The protein and the peptide are represented as cartoons in rainbow color, interacting residues are depicted as lines whereas ligands can be seen as sticks with carbons in blue. Yellow dashes indicate H-bonds.

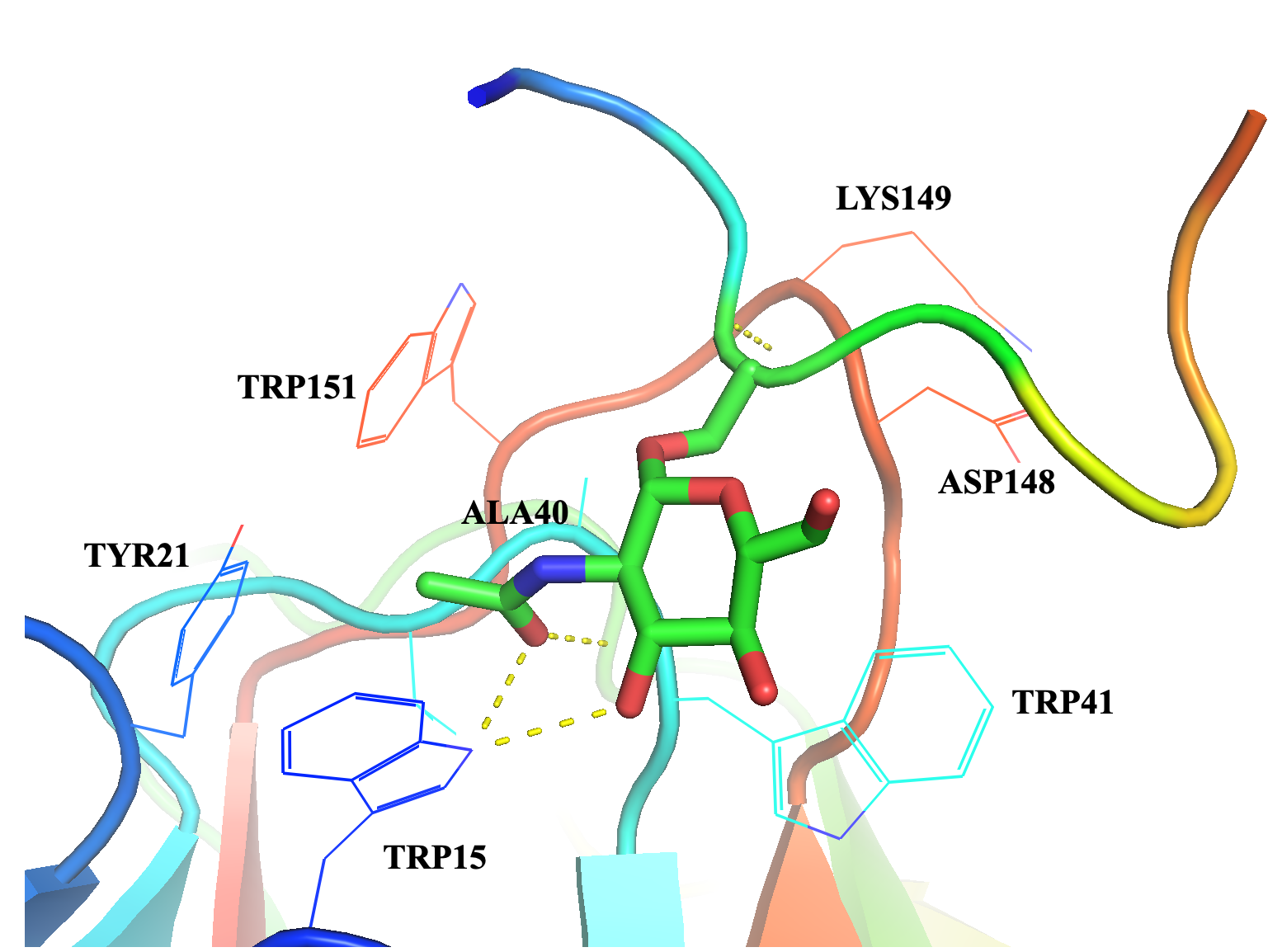
