## Supplementary Figure 6 for "The crystal structure of Nictaba reveals its carbohydrate-binding properties and a new lectin dimerization mode"

**Supplementary Figure 6.** Representative glycan recognition pattern of Nictaba according to glycan array data.

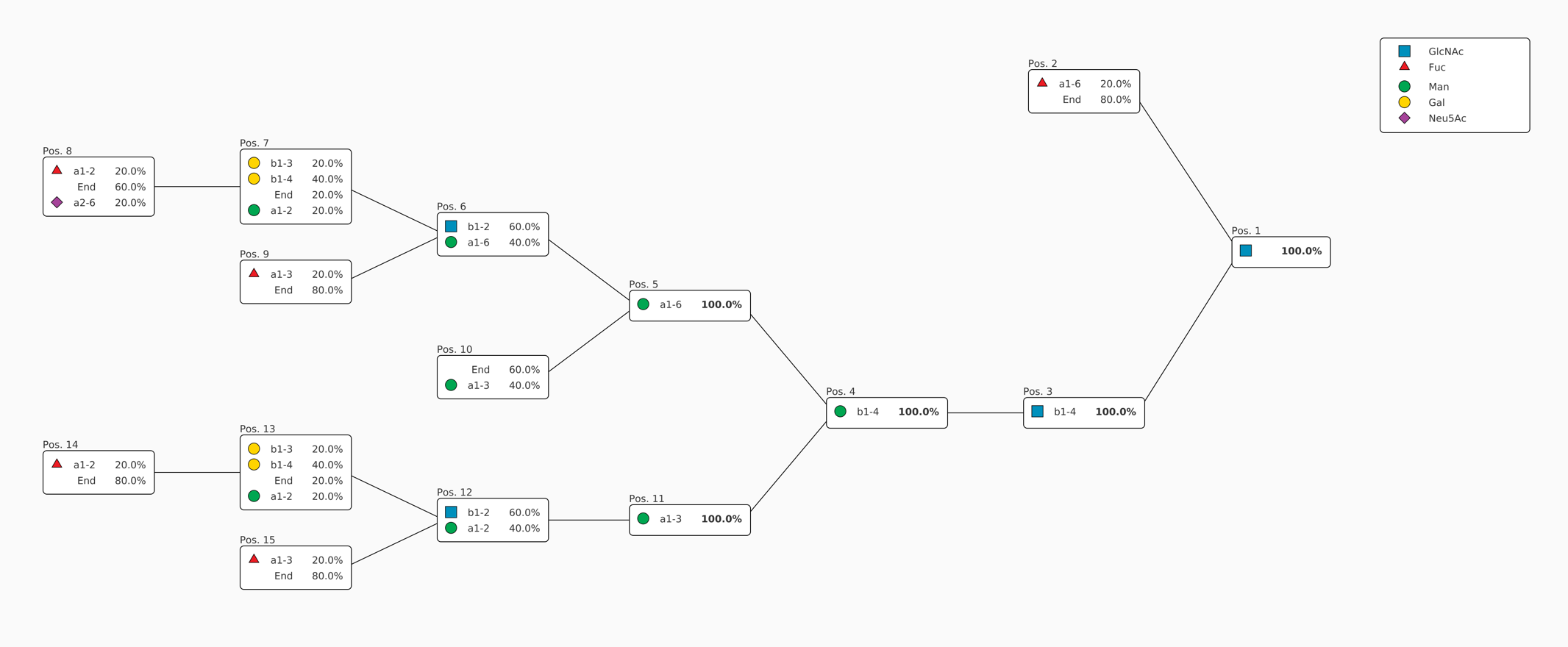
